# The link between static and dynamic brain functional network connectivity and genetic risk of Alzheimer’s disease

**DOI:** 10.1101/2021.03.27.437354

**Authors:** Mohammad S. E. Sendi, Elaheh Zendehrouh, Charles A. Ellis, Jiayu Chen, Robyn L. Miller, Elizabeth Mormino, David H. Salat, Vince D. Calhoun

**Author notes:** Corresponding Author: Mohammad. S. E. Sendi, Vince D. Calhoun.

## Abstract

Apolipoprotein E (APOE) polymorphic alleles are genetic factors associated with Alzheimer’s disease (AD) risk. Although previous studies have explored the link between AD genetic risk and static functional network connectivity (sFNC), to the best of our knowledge, no previous studies have evaluated the association between dynamic FNC (dFNC) and AD genetic risk. Here, we examined the link between sFNC, dFNC, and AD genetic risk with a reproducible, data-driven approach. We used rs-fMRI, demographic, and APOE data from cognitively normal individuals (N=894) between 42 to 95 years of age (mean = 70 years). We divided individuals into low, moderate, and high-risk groups. Using Pearson correlation, we calculated sFNC across seven brain networks. We also calculated dFNC with a sliding window and Pearson correlation. The dFNC windows were partitioned into three distinct states with k-means clustering. Next, we calculated the amount of time each subject spent in each state, called occupancy rate or OCR. We compared both sFNC and OCR, estimated from dFNC, across individuals with different genetic risk and found that both sFNC and dFNC are related to AD genetic risk. We found that higher AD risk reduces within-visual sensory network (VSN) sFNC and that individuals with higher AD risk spend more time in a state with lower within-VSN dFNC. Additionally, we found that AD genetic risk affects whole-brain sFNC and dFNC in women but not in men. In conclusion, we presented novel insights into the links between sFNC, dFNC, and AD genetic risk.

## 1. Introduction

Alzheimer’s disease (AD) is the most prevalent age-related dementia in individuals above 65 years in age (Masters et al., 2015). While global biomedical research efforts for AD prevention have expanded, the number of individuals affected by AD is still growing significantly every year. Even though there is no effective AD therapy to date, some medications can slow down disease progression (Yiannopoulou and Papageorgiou, 2020). It has been hypothesized that AD progression affects brain functional connectivity beginning many years prior to disease onset (Agosta et al., 2012; Demirtaş et al., 2017). As such, knowing how AD risk alters brain connectivity in cognitively normal individuals might shed light on the mechanisms associated with AD development later in life.

While previous studies showed that environmental factors such as diet, living in rural versus urban areas, smoking, not exercising, and infections are risk factors of AD, genetic factors are believed to contribute 70% to AD risk (Elsheikh et al., 2020; Van Cauwenberghe et al., 2016). Apolipoprotein E polymorphic alleles are genetic factors linked to Alzheimer’s disease (AD). There are three common alleles including ε2, ε3, and ε4 that can produce six genotypes such as ε2/ε2, ε2/ε3, ε2/ε4, ε3/ε3, ε3/ε4, and ε4/ε4 in which one gene is inherited from the father and the other is from the mother (Yamazaki et al., 2019). Individuals carrying the ε4 allele have the highest risk of AD and younger mean age at dementia onset compared to those carrying the ε3 and ε2 alleles, whereas individuals with the ε2 allele have the lowest risk and older mean age of dementia onset (Bird, 2008).

Previous studies explored the link between AD’s genetic risk and static functional network connectivity or sFNC (Axelrud et al., 2019; Chiesa et al., 2017; McKenna et al., 2016; Turney et al., 2020). For example, a recent study found that individuals carrying ε4 have lower temporal default mode network (DMN) functional connectivity than those without ε4 (Turney et al., 2020). Another study showed an increase in functional connectivity between the hippocampus and prefrontal/parietal/temporal cortex in healthy individuals carrying ε4 (Zheng et al., 2018). Despite extensive research on the effect of ε4 on rs-fMRI sFNC, the effects of other alleles (i.e., ε2 and ε3) on sFNC have not been explored. Additionally, previous literature that studied the APOE effect on FNC assumed that FNC is static over time and ignored its dynamics. However, recent studies have shown that FNC is highly dynamic during both task and resting conditions (Allen et al., 2014; Mohammad S E Sendi et al., 2021; Mohammad S. E. Sendi et al., 2021). Therefore, we hypothesized here that studying the dynamics of whole-brain FNC might give insight into how the APOE could disrupt FNC in AD.

This study aimed to explore how dFNC and sFNC differ among individuals with different genetic risk for AD using a relatively large dataset (N>850). To model the dFNC of each participant, we utilized a sliding window approach followed by k-means clustering to estimate a set of connectivity states (Allen et al., 2014). Next, we modeled the temporal changes by calculating the occupancy rate (OCR) of each state from the dFNC. Next, we explored the difference between cognitively normal participants with different AD risk levels via statistical analysis on the estimated OCR features and the number of the between-state transitions. In addition, we compared sFNC cell-wise differences between cognitively normal individuals differing in genetic risk of AD.

## 2. Methods and Materials

### 2.1. Participants and dataset

Neuroimaging data of 894 cognitively normal brains (362 females) and their associated demographic information from the longitudinal Open Access Series of Imaging Studies (OASIS)-3 cohort was used in this study (LaMontagne et al., 2019). The participants’ cognitive functionality at the time of scanning was evaluated by the clinical dementia rating scale (CDR) scores, and the CDR scores were equal to 0. The participants’ age at scanning time ranged from 42.46 years to 95.39 years, with a mean of 70.13 years. We divided the data into three groups, including the low genetic risk of AD or LGR_AD (N=135, 63 females), consisting of all individuals with ε2 allele (i.e., ε2/ε2 and ε2/ε3), moderate genetic risk of AD or MGR_AD (N=558, 219 females), containing all individuals with ε3 allele (i.e., ε3/ε3), and high genetic risk of AD or HGR_AD (N=201, 80 females) consisting of all individuals with ε4 allele (i.e., ε3/ε4 and ε4/ε4). The demographic and clinical information of each group is shown in Table 1. No significant age, gender, and mini-mental state examination differences were observed between any pair of groups (p>0.05).

**Table 1.** Demographic and clinical information of subjects

|  | ***LGR-AD*** | ***MGR-AD*** | ***HGR-AD*** |
| --- | --- | --- | --- |
| ***Number*** | 135 | 558 | 201 |
| ***Age*** | 71.22±8.42 | 70.14±8.58 | 69.37±8.59 |
| ***Gender (M/F)*** | 72/63 | 339/219 | 121/80 |
| ***MMSE*** | 28.98±1.17 | 29.15±1.11 | 29.12±1.30 |

**Note:** LGR-AD: Low genetic risk for AD, MGR-AD: Moderate genetic risk for AD, HGR-AD: High genetic risk for AD, MMSE: mini-mental state examination.

### 2.2. Imaging acquisition protocol

High-resolution T2*-weighted functional images were collected by echoplanar (EP) imaging using Trio 3T scanners with 20-channel head coils (Siemens Medical Solutions USA, Inc). The data collecting protocol includes TE =27 ms, TR = 2.2 s, flip angle = 90°, slice thickness = 4mm, slice gap = 4 mm, matrix size = 64, and voxel size of 1 mm × 1 mm × 1.25 mm. The duration of the scanning was 6 minutes.

### 2.3. Preprocessing

Fig. 1 shows the analytic pipeline that we used in this study. The following steps describe the details of our method. In the first step (Step1 in Fig. 1), the first five dummy scans were removed before the preprocessing. We used statistical parametric mapping (SPM12, https://www.fil.ion.ucl.ac.uk/spm/) default slice timing routines. In this method, we used the slice acquired in the middle of the sequence as the reference slice. We applied rigid body motion correction to adjust for participant head movement. Next, we normalized the imaging data to the standard Montreal Neurological Institute (MNI) space using the echo-planar imaging (EPI) template. Finally, we smoothed the images by applying a Gaussian kernel with a full width at half maximum (FWHM) of 6mm.

**Fig. 1.**
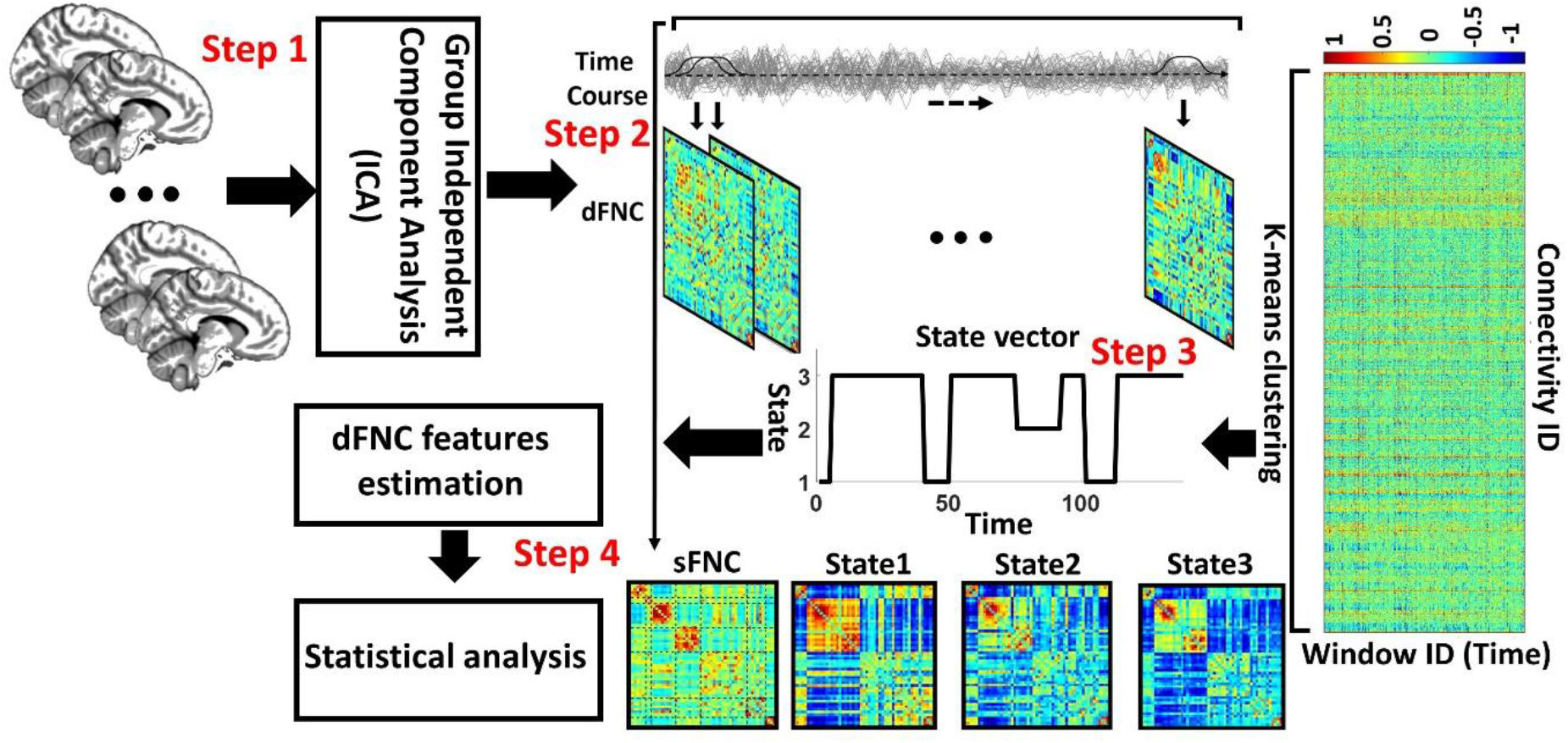
Analytic pipeline: Fifty-three time-course components of the whole brain were identified using group independent component analysis (ICA). In Step2, a taper sliding window was used to segment the time-course signals and calculate the functional network connectivity (FNC). After vectorizing the FNC matrixes, we concatenated them and used k-means clustering, k=3, to group them into three distinct states (Step3). The elbow criteria was used to find the optimal k. In addition, the correlation distance metric was used for the clustering. Then, based on the state vector of each subject, the occupancy rate or OCR features, in total 3 features, were calculated from the state vector of each subject. We next compared the OCR among groups with a two-sample t-test. We adjusted all p values by the Benjamini-Hochberg false discovery rate (FDR) correction in each analysis (Step4).

To obtain comparable components from the dataset in this study, we used a set of robust network priors extracted through the Neuromark pipeline (Du et al., 2020). Group independent component analysis (ICA) was applied on two healthy controls datasets from the human connectome project (HCP: https://www.humanconnectome.org/study/hcp-young-adult/document/1200-individuals-data-release, 823 individuals after the subject selection) and genomics superstruct project (GSP: https://dataverse.harvard.edu/dataverse/GSP, 1005 individuals after the subject selection) to generate the network priors. Fifty-three intrinsic components (ICs) were extracted and arranged in seven functional domains based on anatomic and functional prior knowledge. These seven functional domains were the subcortical network (SCN), auditory network (ADN), sensorimotor network (SMN), visual network (VSN), cognitive control network (CCN), default-mode network (DMN), and cerebellar network (CBN). More details on the extracted ICs are provided in (Mohammad S. E. Sendi et al., 2021).

### 2.4. Dynamic and static functional network connectivity estimation

The sliding window approach with a tapered window (size of 44s and standard deviation of 3 s) was used to estimate the whole-brain dFNC. Pearson correlation was used to estimate the dFNC in each window. With 53 ICs, we had 1378 connectivity features. We concatenated the dFNC estimates of each window for each individual to form a (C × C × T) array (where C=53 denotes the number of ICNs and T=139), which represented the changes in brain connectivity between ICs as a function of time (Step 2 in Figure 1).

### 2.5. Dynamic functional network connectivity clustering

We separated the windowed FNCs into a set of clusters (or states) with k-means clustering. The optimal number of clusters or k was set to three based upon the elbow criterion, e.g., “the ratio of within-cluster to between cluster distance. Pearson correlation was used as a distance metric, and 1000 iterations were used. The output of k-means clustering was three states for each group and a state vector for each individual. The state vector represents changes in wholebrain FNC over time. Next, we calculated the proportion of each participant’s time in each state, called occupancy rate (OCR) hereafter. Having three states, we estimated three OCRs for each individual.

### 2.6. Statistical analysis

A two-sample t-test was used to compare the OCR (number of null hypotheses or N=3) of each pair of groups. Similarly, a two-sample t-test was used to compare the sFNC (N=1387) of each pair of groups. We adjusted all p values with Benjamini-Hochberg false discovery rate (FDR) correction in both OCR and sFNC analysis (Yoav Benjamini; Yosef Hochberg, 1995).

## 3. Results

### 3.1. Genetics risk associated with sFNC

The average sFNC of each group is shown in Fig. 2A. Also, Fig.2B shows the cell-wise FNC differences between each pair of groups. We used a two-sample t-test to find the differences between groups. Significant group differences that passed the multiple comparison tests are shown with an asterisk (FDR corrected q=0.05). In this figure, the red and blue colors show the positive and negative difference, respectively. Fig. 2B (left panel) shows the cell-wise difference between LGR_AD and MGR_AD (MGR_AD-LGR_AD). We did not observe a significant sFNC difference between LGR_AD and MGR_AD groups after FDR correction. While the difference between LGR_AD and HDR_AD (or HGR_AD-LGR_AD) was significant in some networks, as shown in Fig. 2B (middle panel). However, the pattern was widespread across the whole brain.

**Fig. 2.**
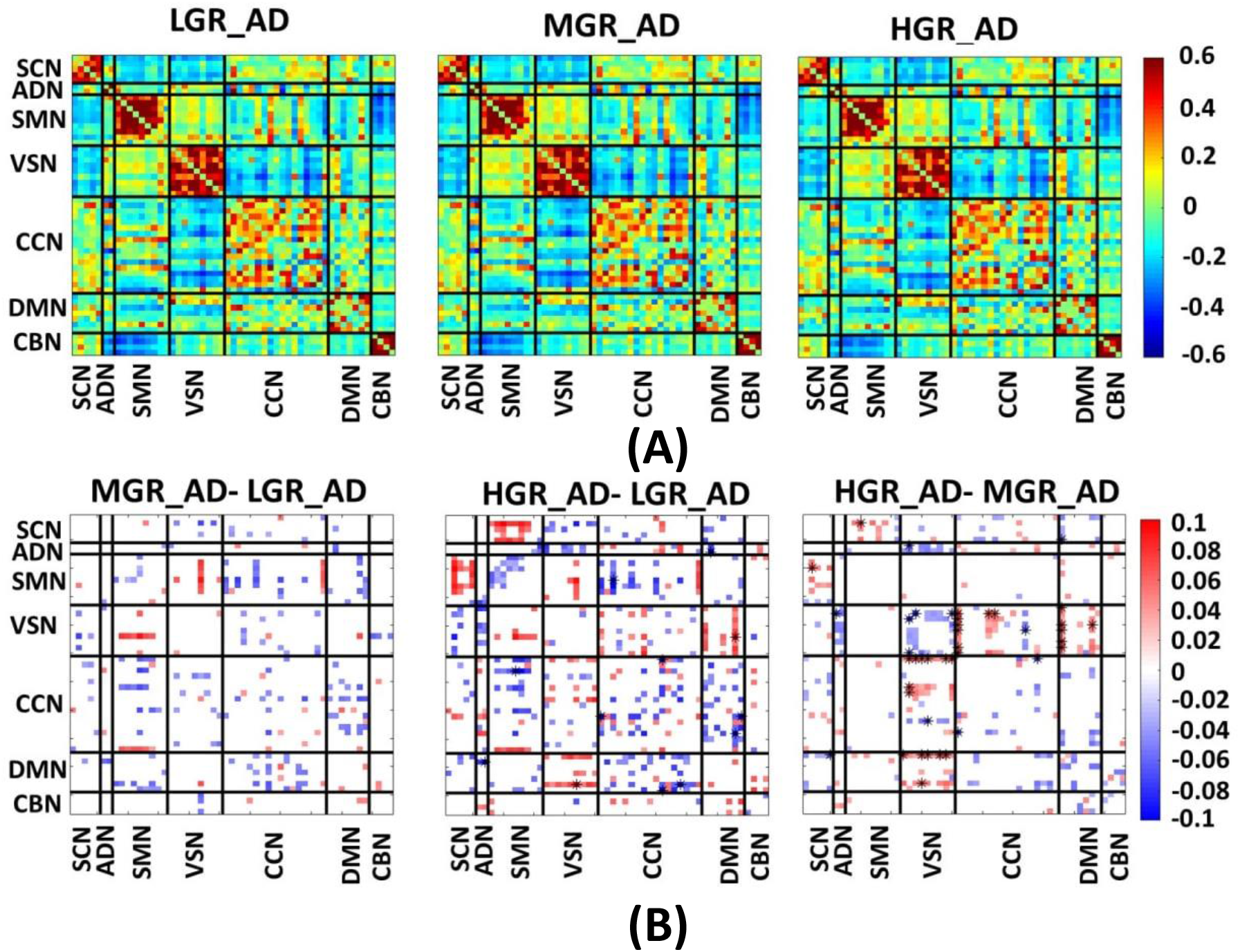
Estimated sFNC for each group. **A)** Estimated sFNC for LGR-AD (left panel), MGR-AD (middle panel), and HGR-AD (right panel). **B)** The sFNC difference between each pair of groups. Significant group differences passing the multiple comparison threshold are marked by asterisks (false discovery rate [FDR] corrected, q = 0.05). The colorbar shows the intensity of sFNC values. SCN: subcortical network SCN, ADN: auditory network, SMN: sensory motor network, VSN: visual sensory network, CCN: cognitive control network, DMN: default mode network, and CBN: cerebellar network. LGR-AD: Low genetic risk of AD, MGR-AD: Moderate genetic risk of AD, HGR-AD: High genetic risk of AD

In contrast, the cell-wise FNC difference between MGR_AD and HDR_AD (or HGR_AD-MGR_AD) was more focused on VSN, as shown in Fig. 2B (right panel). As shown in this figure, we found that individuals with a higher risk of AD have less VSN connectivity than those with an intermediate risk of AD. In comparison, the connectivity between VSN and CCN and between VNS and DMN was higher for individuals with higher AD risk.

### 3.2. Dynamic functional network connectivity states

Fig.3 shows three distinct dFNC states estimated by k-means clustering. State1 and state 2 show more positive connectivity within CCN, within CBN, within SMN, and within VSN compared with state3. State 2 offers the most positive connectivity among sensory domains (i.e., ADN, SMN, and VSN). Meanwhile, the connectivity between these three domains with the rest of the brain is negative in this state. Additionally, the connectivity between SMN and the rest of the brain is relatively high in state1.

**Fig. 3.**
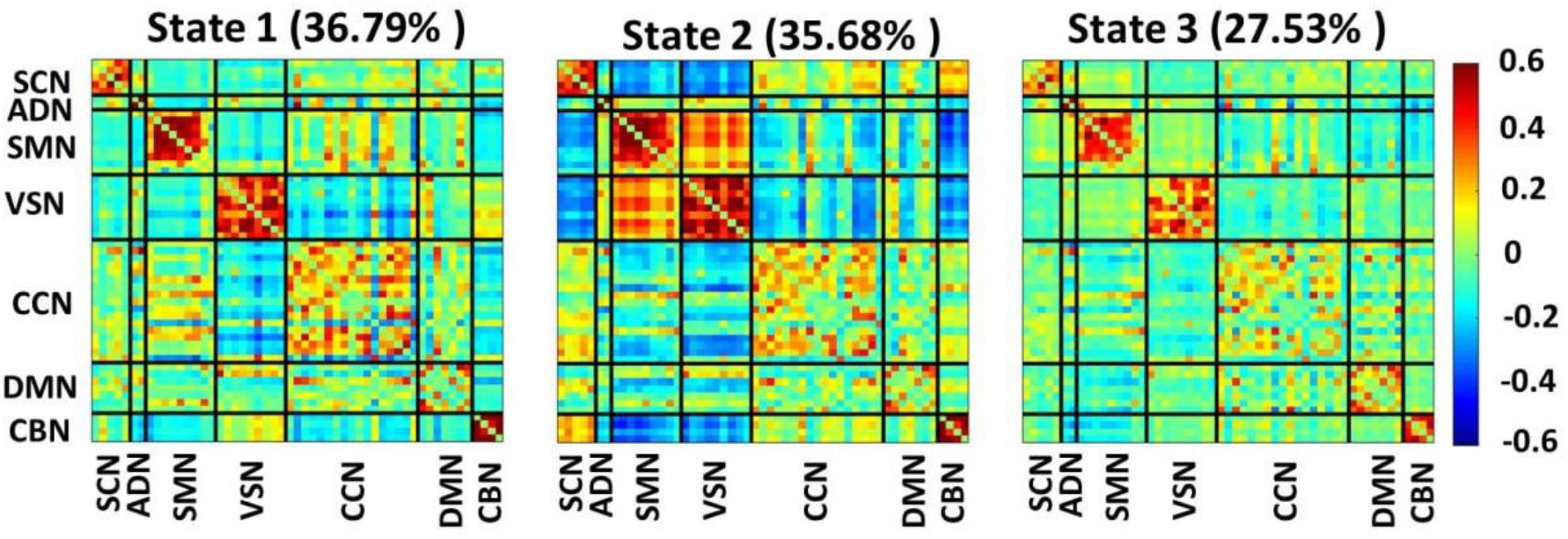
Three dFNC states identified with the k-means clustering method. Each state is a 53×53 matrix in which positive connectivity is shown with red, and the negative connectivity is shown in blue. We put all 53 components in 7 domains including the subcortical network (SCN), auditory network (ADN), sensory motor network (SMN), visual sensory network (VSN), cognitive control network (CCN), default mode network (DMN), and cerebellar network (CBN). We found individuals spent 36.79 %, 35.68 %, and 27.53 % in state 1, state 2, and state 3, respectively.

### 3.3. Genetics risk associated with dFNC features

We compared the OCR of each state across three groups of individuals. The results are shown in Fig. 4A. As this figure shows, we found that the OCR of state1 was significantly less for HGR_AD than that of LGR_AD. In contrast, HGR_AD had higher OCR than LGR_AD in state 3. Additionally, we did not find any significant OCR difference across groups in state 2. Besides, no significant difference was observed between MGR_AD and LGR_AD and between HGR_AD and MGR_AD.

**Fig. 4.**
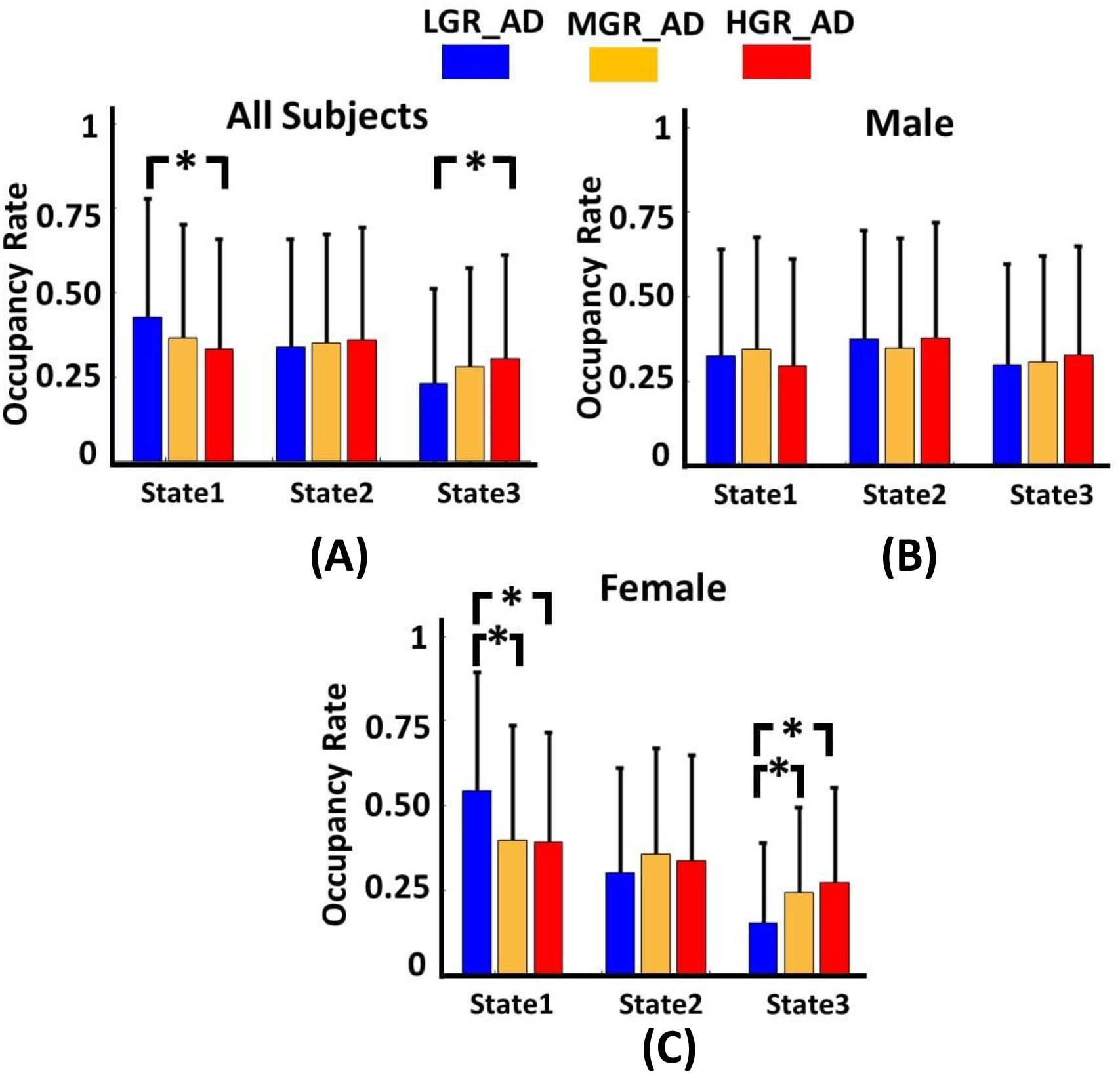
Genetic risk effects on dFNC features. **A)** All individuals: The occupancy rate (OCR) of each group in state 1, state 2, and state 3. The individuals with ε2 allele spent more time in state 1 than those individuals without the ε2 allele (corrected p<0.05). **B)** Women: The occupancy rate (OCR) of each group in state 1, state 2, and state 3. The individuals without the ε2 allele spent more time in state 3 than those individuals with the ε2 allele (corrected p<0.05). **C)** Men: The occupancy rate (OCR) of each group in state 1, state 2, and state 3. No significant difference was observed among groups. LGR-AD: Low genetic risk of AD, MGR-AD: Moderate genetic risk of AD, HGR-AD: High genetic risk of AD

### 3.4. Gender effect on sFNC and dFNC features

To consider sex effects on the results, we separated men and women in each group of individuals and repeated our analysis. The group sFNC difference for each sex is shown in Fig 5. Fig. 5 A shows the sFNC difference for each male participant group, and Fig. 5B shows similar results for women. Interestingly, we did not find any significant sFNC difference across groups for males. In contrast, the sFNC difference between LGR_AD and HGR_AD was significant for the women, as shown in Fig. 5B (left panel). We did not observe a significant difference between women LGR_AD and MGR_AD and between women MGR_AD and HGR_AD.

**Fig. 5.**
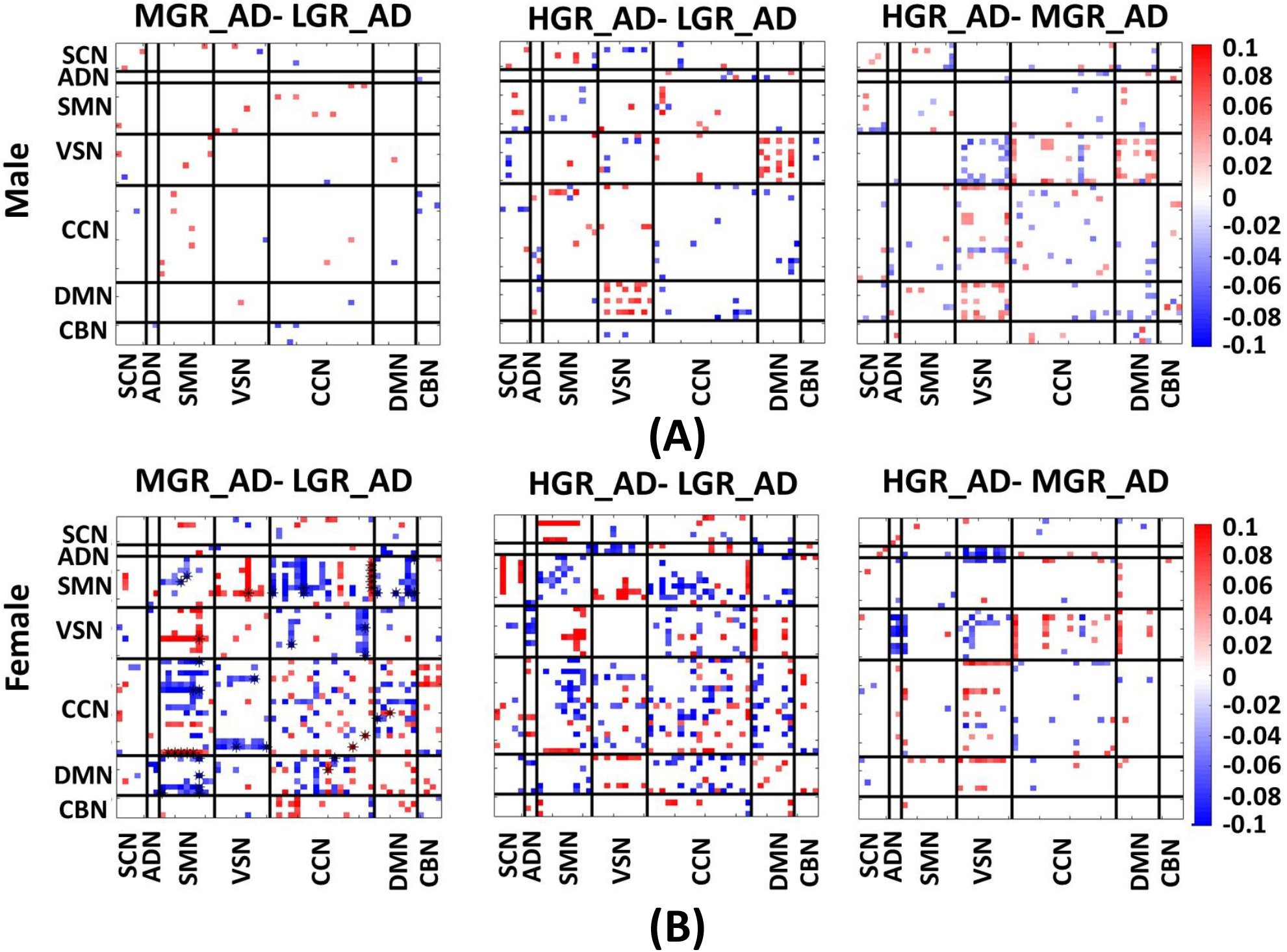
Sex effects on sFNC. **A)** sFNC differences between pairs of groups for men. **B)** sFNC difference between pairs of groups for women. Significant group differences passing the multiple comparison threshold are marked by asterisks (false discovery rate [FDR] corrected, q = 0.05). The colorbar shows the intensity of sFNC values. SCN: subcortical network SCN, ADN: auditory network, SMN: sensory motor network, VSN: visual sensory network, CCN: cognitive control network, DMN: default mode network, and CBN: cerebellar network. LGR-AD: Low genetic risk of AD, MGR-AD: Moderate genetic risk of AD, HGR-AD: High genetic risk of AD

We also studied group differences in dFNC features for both men and women separately. We did not observe a significant group difference in the OCR of each state (Fig. 4B) for men individuals like sFNC result. In comparison, the group difference was significant for the women. In more detail, we found a significant OCR difference between LGR_AD and MGR_AD and between LGT_AD and HGR_AD in state 1 and state 3. While sFNC features did not differentiate between these groups.

## 4. Discussion

Previous studies showed that FNC estimated from rs-fMRI is highly dynamic even without external input (Allen et al., 2014; Mohammad S. E. Sendi et al., 2021). Therefore, here we hypothesized that the genetic risk of AD not only alters the strength of the functional connectivity between pairs of brain networks as would be shown in sFNC but also the dynamic fluctuations of connectivity among those networks as would be shown in dFNC. To the best of our knowledge, the present study is the first to report a link between AD genetic risk and dFNC estimated from rs-fMRI recorded of cognitively normal participants. We also compared the results obtained from both sFNC and dFNC data as the measures are complementary and could provide distinct insights into the relationship between brain network connectivity and the genetic risk of AD. Lastly, we examined the effects of an individual’s sex on the degree to which sFNC, dFNC, and genetic risk of AD are associated.

We used a large dataset and data-driven, reproducible methods in our study. We used 6-minute sessions of rs-fMRI data from 894 cognitively normal individuals with different AD genetic risks and put them into three groups, including individuals carrying at least one ε2 (i.e., ε2/ε2 and ε2/ε3), individuals carrying only ε3 (i.e., ε3/ε3), and individuals carrying at least one ε4 (i.e., ε3/ε4 and ε4/ε4). We used a sliding window approach to calculate the whole-brain dFNC over time and k-means clustering to partition the whole-brain dFNC into three distinct states. Next, we compared the OCR for each state among individuals with differing in genetic risk for AD. We also compared sFNC across individuals with different risks of AD.

We observed a widespread effect of genetic risk on the sFNC of cognitively normal individuals, which is consistent with previous studies (Sheline et al., 2010; Trachtenberg et al., 2012). We found that genetic risk affects the VSN significantly more than other brain networks. In more detail, we found that individuals carrying ε4 have less within-VSN functional connectivity compared to those individuals carrying ε3 and not ε2. A decrease in VSN activity during a visual task has been reported in individuals carrying ε4 (Smith et al., 1999). A recent study reported functional connectivity decreases in the primary, secondary, and associative visual cortices for the cognitively normal individuals with ε4 allele compared with ε4 allele non-carriers (McKenna et al., 2016). We also found that cognitively normal individuals carrying ε4 have higher VSN/CCN than individuals with moderate AD risk and more increased VSN/DMN connectivity relative to individuals with moderate and lower AD risk. We hypothesize that this higher VSN/CCN connectivity in ε4 carriers could be a compensatory mechanism to offset the VSN FC reduction in these individuals. A similar compensatory mechanism in functional connectivity has been reported in individuals with ε4, enabling them to achieve the same performance level as individuals with ε3 in a memory task (Bondi et al., 2005). It is worth mentioning that we did not find a significant sFNC difference between the individuals with the lowest risk and the moderate risk of AD, which suggests that a modest risk of AD does not change the brain FNC to the extent that the highest risk of AD does.

We also explored the whole-brain dFNC across cognitively normal individuals with different genetic risks for AD. We observed a significant difference in the OCR of individuals carrying ε4 and individuals having ε2. At the same time, we did not observe any significant differences between those with ε2 and ε3 and those with ε3 and ε4. In more detail, we found that individuals carrying ε4 spend more time than those carrying ε2 in state3 with lower within-VSN functional connectivity (relative to state1 and state2). This result is consistent with the result obtained from the sFNC data, which showed that individuals with higher AD genetic risk have less within-VSN functional connectivity. That supported our hypothesis about the genetic risk effects on both the strength and the temporal pattern of FNC estimated from rs-fMRI.

Additionally, we found that individuals with ε4 spend more time in state3 with lower SMN, lower CCN, and lower CBN functional connectivity (relative to state1 and state2), while we did not observe a significant difference in those networks by analyzing sFNC data. These results might reveal some new evidence of the CCN, VSN, and CBN’s role in differentiating individuals with different genetic risk for AD. We did not observe a significant difference in OCRs of the individuals with a moderate and lower risk of AD, consistent with the sFNC analysis results. These pieces of evidence might suggest that a moderate genetic risk of AD does not affect either brain sFNC or dFNC.

We observed a considerable difference in sFNC when comparing the individuals having ε3 with those having ε4. However, no significant difference between the two groups was observed by looking at OCR. While previous studies only analyzed sFNC (Axelrud et al., 2019; Chiesa et al., 2017; McKenna et al., 2016; Turney et al., 2020), the results reported above demonstrated the importance of examining both sFNC and dFNC data to differentiate individuals with different AD risks.

We also separated men and women to examine the effect of sex on our results. We did not observe a significant difference across these three groups of men for either sFNC or dFNC data. While a significant difference between women carrying ε4 and ε2 and between women carrying ε4 and ε3 was observed in the dFNC analysis, we only found a significant difference between the women with ε2 and ε4 in the sFNC analysis. We did not observe a significant difference between individuals having ε3 and ε4 in either analysis. A recent study analyzed 5496 healthy individuals carrying ε4 and showed that AD’s conversion rate is significantly higher for women (Altmann et al., 2014). Our current research shows that sex differences significantly contribute to differentiating cognitively normal individuals with different AD genetic risks based on their rs-fMRI, which might explain the study above’s finding. Although previous studies considered the ε3 allele the be a neutral factor in AD, a recent study claimed that the ε3 allele might be a protective factor rather than a neutral one (de-Almada et al., 2012). Our results may suggest that the ε3 allele is a more significant risk factor for AD in women than in men.

It should be noted that our study does have some limitations. In particular, previous studies have indicated that risks in addition to genetic risk can lead to AD (Durazzo et al., 2014; Rahman et al., 2020; Thomas et al., 2020). For example, a recent study showed that individuals with diabetes and the ε4 allele demonstrated a faster functional decline than those without diabetes (Thomas et al., 2020). Other confounding factors like smoking (Durazzo et al., 2014), physical activity (Meng et al., 2020), and education levels (Meng and Arcy, 2012) could introduce some bias into our results. This information was not included in the dataset. Future studies are needed to explore AD genetic risk factors combined with other potential risk factors in both sFNC and dFNC data.

In conclusion, by analyzing the link between AD genetic risk with sFNC and dFNC for the first time, we found that AD genetic risk affects both sFNC and dFNC and that each analysis provides information about different aspects of the effects of AD risk on brain connectivity. When analyzing sFNC data, it was possible to differentiate people with lower risk from those with higher risk and people with moderate risk from those with higher risk. An analysis of dFNC showed that people with low risk could be discriminated from those with higher risk and that the SMN and CBN helped differentiate the two groups. When analyzing sFNC and dFNC from individuals of both sexes, we found that a higher risk of AD is associated with a reduction in within-VSN connectivity and an increase in VSN/DMN connectivity, potentially as a compensatory mechanism. However, when analyzing only women, we did not observe a similar compensatory mechanism. The lack of a compensatory mechanism in women could explain the higher AD conversion rate in women that has been identified in previous studies (Altmann et al., 2014). Additionally, our findings suggested that having only an ε3 allele could be more of a risk factor in women than in men. Our results shed new light on the genetic risk interactions for AD and brain connectivity in cognitively normal individuals and could assist future diagnostic and treatment efforts.

## 5. Acknowledgments

We thank who collected the data and the participants of this study.

## 8. Author’s contributions

Mohammad S. E. Sendi developed the study, conducted data analysis, interpreted the results, and wrote the original manuscript draft. Elaheh Zendehrouh conducted data analysis. Charles A. Ellis wrote the original manuscript draft. Jiayu Chen provided critical review and edited the manuscript. Robyn L. Miller provided critical review and edited the manuscript. Elizabeth Mormino provided critical review and edited the manuscript. David H. Salat provided critical review and edited the manuscript. Vince D. Calhoun developed the study, interpreted the results, edited the original draft, and provided critical review to the initial draft. All authors approved the final manuscript.

